# Deep Learning-Guided Design of Candidate Synthetic Cis-Regulatory Elements for a TCF7+ Progenitor-Exhausted-like CD8+ T-Cell State

**DOI:** 10.64898/2026.11.23.768730

**Authors:** Nasreen Buhn, Matthew Lee, Dokyoon Kim, Saar Gill

**Affiliations:** Center for Cellular Immunotherapies, Perelman School of Medicine, University of Pennsylvania, Philadelphia, PA, USA; Institute for Immunology & Immune Health (I3H), Perelman School of Medicine, University of Pennsylvania, Philadelphia, PA, USA; Department of Biostatistics, Epidemiology and Informatics, Perelman School of Medicine, University of Pennsylvania, Philadelphia, PA, USA; Department of Systems Pharmacology and Translational Therapeutics, Perelman School of Medicine, University of Pennsylvania, Philadelphia, PA, USA; Institute for Biomedical Informatics, Perelman School of Medicine, University of Pennsylvania, Philadelphia, PA, USA; Division of Hematology-Oncology, Department of Medicine, University of Pennsylvania, Philadelphia, PA, USA; Center for Cell Therapy and Transplant, University of Pennsylvania, Philadelphia, PA, USA; Parker Institute for Cancer Immunotherapy at University of Pennsylvania, University of Pennsylvania, Philadelphia, PA, USA

## Abstract

Progenitor-exhausted CD8+ T cells (Tex-prog) are essential for sustaining T-cell-mediated immunity during chronic infection and cancer and therefore offer a promising immunotherapy target. Synthetic cis-regulatory elements (CREs) could enable selective transgene expression in Tex-prog cells, but whether they can distinguish this population from closely related CD8+ T-cell exhaustion states remains unclear. Here, we integrated publicly available paired single-cell ATAC-seq and single-cell RNA-seq datasets from mouse infection and tumor models and identified an ATAC-defined Tex-prog-like population (C1). We used the CREsted framework to fine-tune a pretrained Borzoi model to predict chromatin accessibility across C1 and ten comparator populations. We then used CREsted to design candidate synthetic CREs with predicted preferential accessibility in C1. The fine-tuned model achieved an average precision of 0.242 for identifying C1-selective regions on chromosomes held out from fine-tuning, approximately tenfold higher than the random-ranking baseline. Sequence optimization generated 196 1,000-bp designs with at least twofold higher predicted accessibility in C1 relative to the strongest competing population across eight DNA backgrounds. Motif enrichment and sequence-attribution analyses identified sequence patterns associated with AP-1, NFAT, POU, EGR, and TCF/LEF transcription factor families. Together, these findings identify candidate synthetic CREs with predicted preferential accessibility in a Tex-prog-like CD8+ T-cell population and provide a foundation for future experimental evaluation of their functional enhancer activity and cell-state specificity.

## Introduction

Persistent antigen exposure during chronic viral infections and cancer redirects CD8+ T-cell differentiation towards an exhaustion trajectory.^1^ Self-renewing progenitor-exhausted CD8+ T cells (Tex-prog) give rise to a transitional intermediate-exhausted cell state (Tex-int), which exhibits increased expression of effector molecules. Tex-prog and Tex-int cells further differentiate into terminally exhausted T cells (Tex-term), a state characterized by decreased and altered effector function and limited proliferative capacity.^2^ As a result, Tex-term cells have a reduced ability to sustain effective immune responses in chronic viral infection and cancer.^3^

PD-1 blockade has emerged as an effective immunotherapy to reinvigorate exhausted CD8+ T cells. However, Tex-term tumor-infiltrating lymphocytes (TILs) exhibit a limited response to PD-1 blockade, and exhausted T cells (Tex) reinvigorated during chronic viral infection can become re-exhausted upon persistent antigen exposure.^1,4^ Although PD-1 blockade can improve CD8+ T-cell function, it does not entirely reverse exhaustion or restore cells to a functional state.^5,6^ Instead, the proliferative response observed following PD-1 blockade arises predominantly from TCF1-expressing Tex-prog-like cells.^5,6^ Furthermore, melanoma patients with a higher proportion of Tex-prog cells have been found to have a longer response duration to PD-1 blockade therapy.^4^ These findings establish Tex-prog cells as a crucial population for maintaining durable immune responses and point to the Tex-prog population as a promising immunotherapy target that may improve the durability and effectiveness of CD8+ T-cell-mediated immunity.

Engineering state-specific transgene expression in CD8+ Tex-prog cells could enable Tex-prog-targeted therapies while limiting expression in other Tex states. Lan et al. identified Tex-prog-associated chromatin remodeling in response to TCR stimulation and a Tex-prog-specific enhancer regulating Slamf6 expression, providing evidence for distinct cis-regulatory activity in Tex-prog cells.^7^ Furthermore, recent studies have demonstrated that regulatory differences can be leveraged to engineer synthetic cis-regulatory elements (CREs). Gosai et al. developed synthetic CREs that drive cell-type-specific gene expression, while Zhang et al. engineered synthetic promoters that respond to T-cell exhaustion.^8,9^ However, it remains unclear whether synthetic CREs could distinguish Tex-prog cells from closely related Tex states.

Deep learning models can learn relationships between DNA sequences and cell-state-specific chromatin accessibility. These models can subsequently be leveraged to guide the design of state-specific CREs.^10,11^ Using publicly available paired mouse single-cell RNA-seq and ATAC-seq data from acute and chronic LCMV infection and tumor models, we identified a Tex-prog-like CD8+ T-cell population (C1). Here, we employ CREsted (cis-regulatory element sequence training, explanation and design), a software package developed by Kempynck et al. for enhancer modeling and sequence design, to generate candidate synthetic CREs predicted to be preferentially accessible in C1.^12^

## Results

### ATAC clustering identifies a distinct Tex-prog-like CD8+ T-cell population

Leiden clustering of Harmony-integrated scATAC-seq data from 111,347 CD8+ T cells that passed quality control identified four clusters at a resolution of 0.3 (Figure 1D). The C1 cluster contained 4,440 cells that grouped together in both the scRNA-seq and scATAC-seq UMAP embeddings (Figures 1D and 2A). C1 was composed primarily of cells from GSE230363 and GSE267584, with the largest proportion from LCMV Clone 13-infected mice (Figures 1A and 2C). Based on transferred Green et al. reference annotations, 61.6% of C1 cells were classified as Tex-prog (Figures 1B and 2C).^13^ Conversely, C1 contained 2,734 of 5,147 Tex-prog cells (53.1%) and 82 of 3,104 TIL Tex-prog cells (2.6%), indicating that C1 is enriched for, but does not encompass, the Tex-prog compartment. In contrast, ImmGenT annotations classified 45.0% of C1 cells as CD8.wX, a provisional population of unclear identity, and 22.7% as CD8.P, a proliferating CD8+ T-cell population (Figures 1C and 2C).^14^

**Figure 1.**
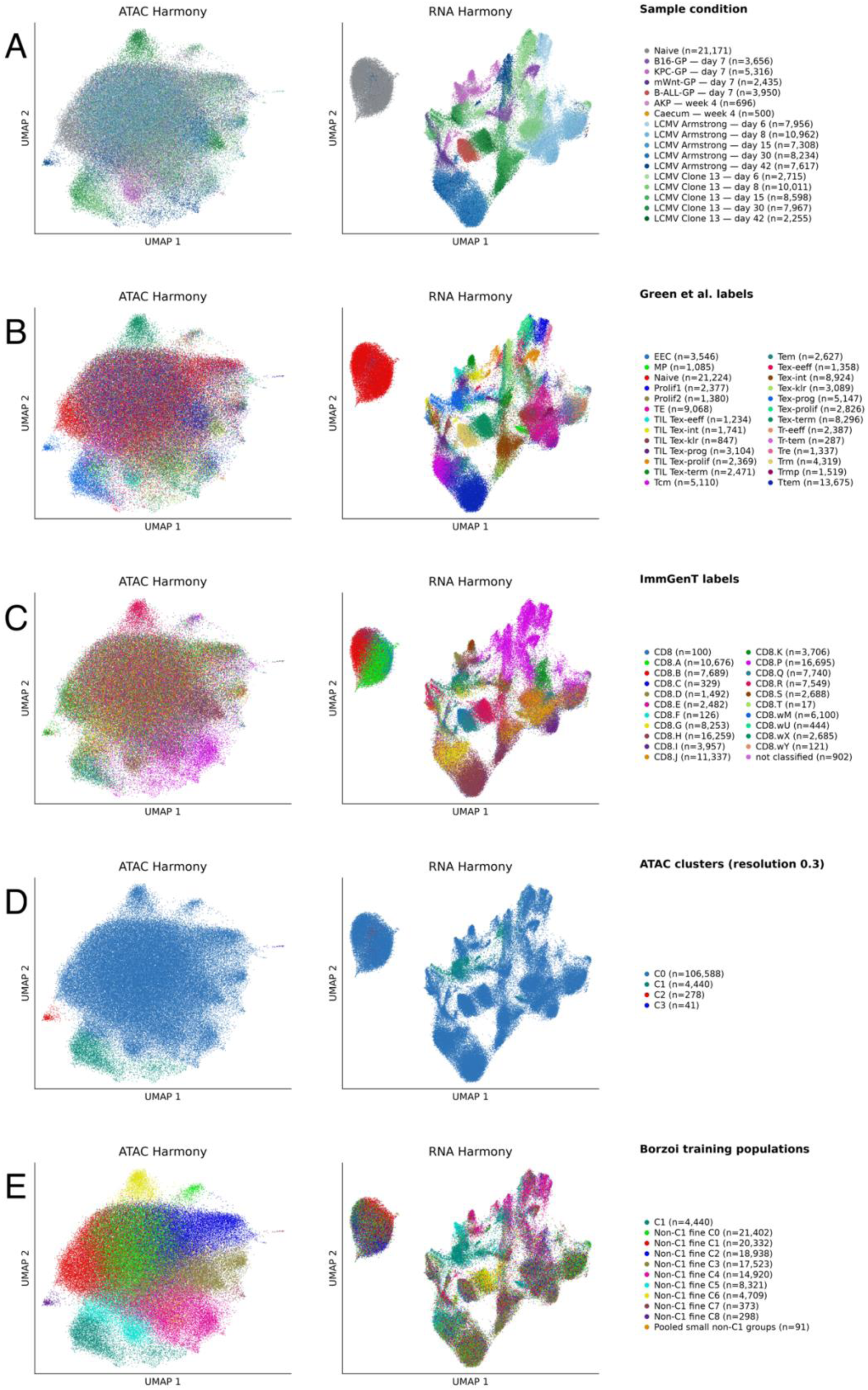
ATAC clustering identifies C1 and comparator populations for Borzoi training. Paired scATAC-seq (left) and scRNA-seq (right) UMAPs of 111,347 CD8+ T cells. Embeddings were generated from Harmony-corrected latent semantic indexing (LSI) components for scATAC-seq and principal components for scRNA-seq. The same cells and UMAP coordinates are maintained across A-E. **A** Cells colored by sample condition and collection time. **B** Cells colored by transferred Green et al. reference annotations. **C** Cells colored by ImmGenT annotations. **D** Cells colored by scATAC-seq-based Leiden clustering at a resolution of 0.3. **E** Cells colored by the 11 populations used for Borzoi training: C1, nine non-C1 groups derived from resolution 1.0 scATAC-based Leiden clustering, and a pooled group of smaller non-C1 populations.

**Figure 2.**
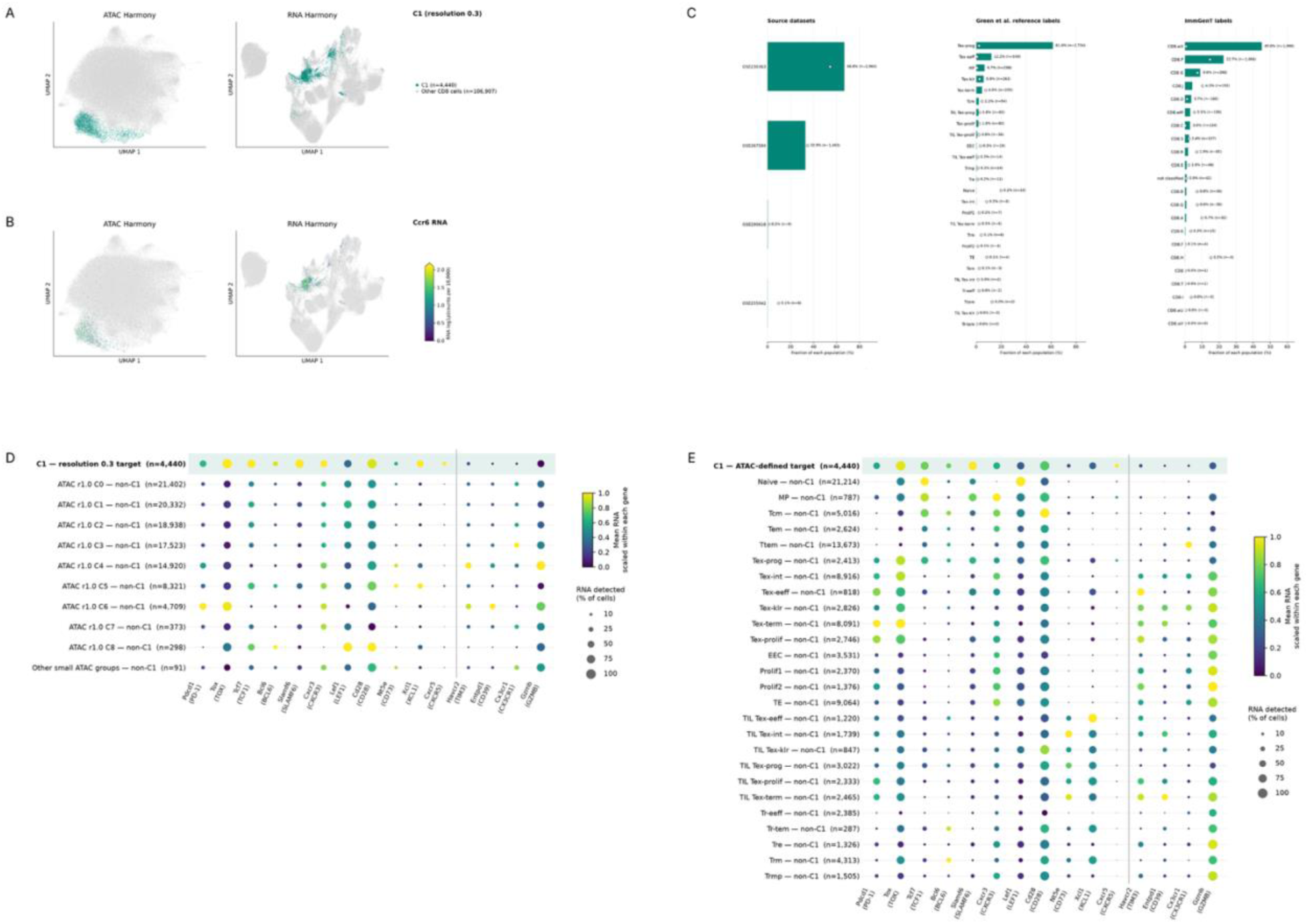
C1 exhibits a Tex-prog-like gene expression profile. A, **B** Paired scATAC-seq (left) and scRNA-seq (right) UMAPs of 111,347 CD8+ T cells. Embeddings were generated from Harmony-corrected latent semantic indexing (LSI) components for scATAC-seq and principal components for scRNA-seq. The same cells and UMAP coordinates are maintained across A and B. **A** The ATAC-defined C1 population. **B** *Ccr6* RNA expression. **C** Composition of C1 by source dataset, transferred Green et al. reference annotation and ImmGenT annotation. Teal bars represent percentages within the C1 population, and open white circles represent the corresponding percentages of non-C1 cells. **D, E** Expression of Tex-prog-defining genes across the 11 Borzoi training populations (D) and Green et al. reference populations excluding C1 cells (E).

Expression of Tex-prog-associated marker genes in the paired scRNA-seq data further supported the identification of C1 as a Tex-prog-like population (Figures 2D and 2E).^2^ *Tcf7* was detected in 70.5% of C1 cells, *Tox* was expressed in 96.8%, *Slamf6* was expressed in 76.9%, and *Cd28* was expressed in 92.9%. In contrast, genes associated with more differentiated exhaustion states were detected in low percentages in C1: *Havcr2* was expressed in 12.9% of C1 cells, *Entpd1* was expressed in 5.6%, and *Cx3cr1* was expressed in 5.9% (Figures 2D and 2E).^2^ C1 exhibited high Cxcr5 expression (Figure 2D). Furthermore, *Ccr6* expression was predominantly concentrated in C1 cells (Figure 2B). Previous work by Giles et al. identified *Ccr6* expression in a Tex-prog population subset.^15^

### Fine-tuned Borzoi learns sequence features associated with C1-selective chromatin accessibility

We fine-tuned the pretrained mouse Borzoi model to predict normalized ATAC fragment counts for C1 and 10 comparator populations from 2,048-bp DNA sequences (Figure 1E). Model training completed after 15 epochs, and the epoch-8 checkpoint was selected based on the highest validation macro average precision (AP) (Figure 3B), although macro-AP varied by less than 0.01 across epochs 7-15. C1-specific validation AP at epoch 8 was 0.262.

**Figure 3.**
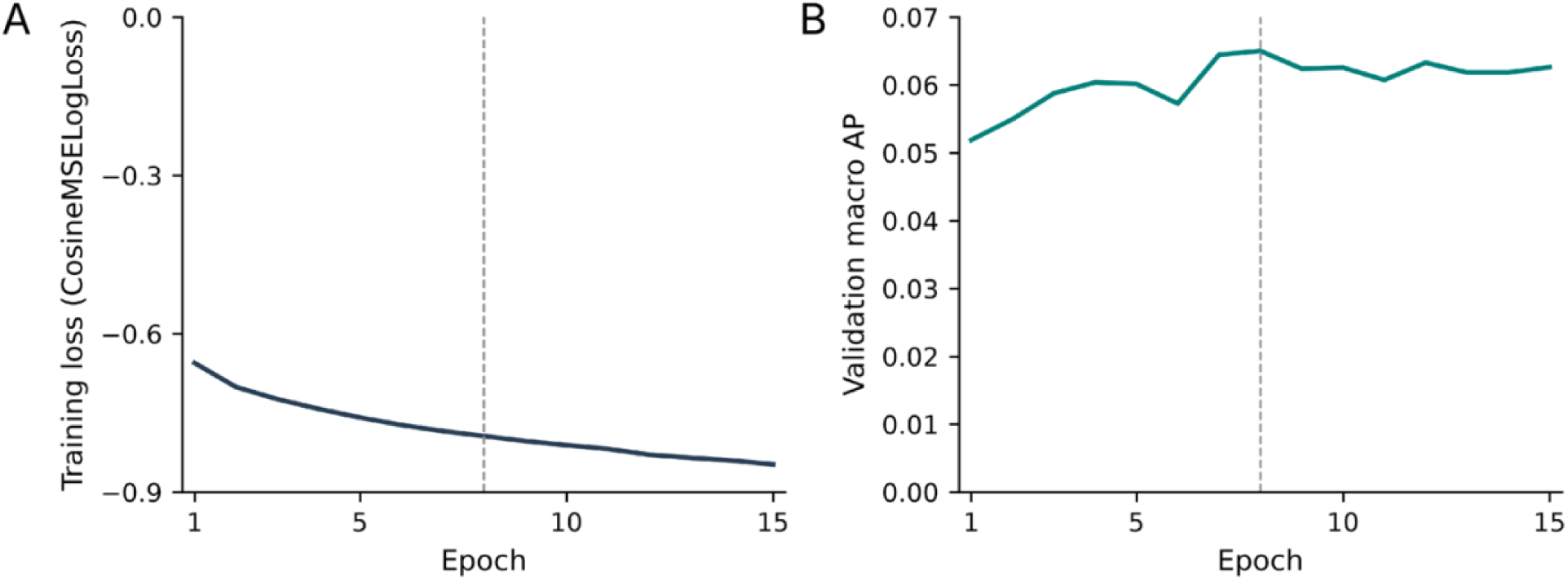
Borzoi validation AP peaked at epoch 8 during fine-tuning. **A** Training loss (CosineMSELogLoss) over 15 epochs. **B** Validation macro-averaged average precision (AP) over 15 epochs. Dashed vertical lines indicate epoch 8.

Among the 37,197 held-out test chromosome regions, 2.35% met our C1-selectivity criteria. Ranking regions by predicted C1 preference resulted in an AP of 0.242, approximately 10-fold above the value of 0.0235 expected by chance at this prevalence. Of the top 20, 50, 100, and 500 ranked regions, 12, 24, 52, and 197 were C1-selective, corresponding to precisions of 60.0%, 48.0%, 52.0%, and 39.4%. Overlap between these test regions and the original Borzoi pretraining regions was not assessed.

When the pooled small population was included as a comparator in both the C1-selective peak definition and the prediction ranking score, 2.14% met the selectivity criteria. C1 AP was 0.229. Of the top 20, 50, 100 and 500 ranked regions, 12, 24, 51 and 182 were C1-selective, corresponding to precisions of 60.0%, 48.0%, 51.0%, and 36.4%, respectively. This sensitivity analysis evaluated the same epoch-8 checkpoint selected using the primary validation criteria.

### Predicted C1 selectivity after optimization of 200-, 500-, and 1,000-bp inserts

Using the frozen epoch-8 checkpoint, 200-bp, 500-bp and 1,000-bp inserts were optimized for separation between C1 and the strongest competing population (Figure 4).

**Figure 4.**
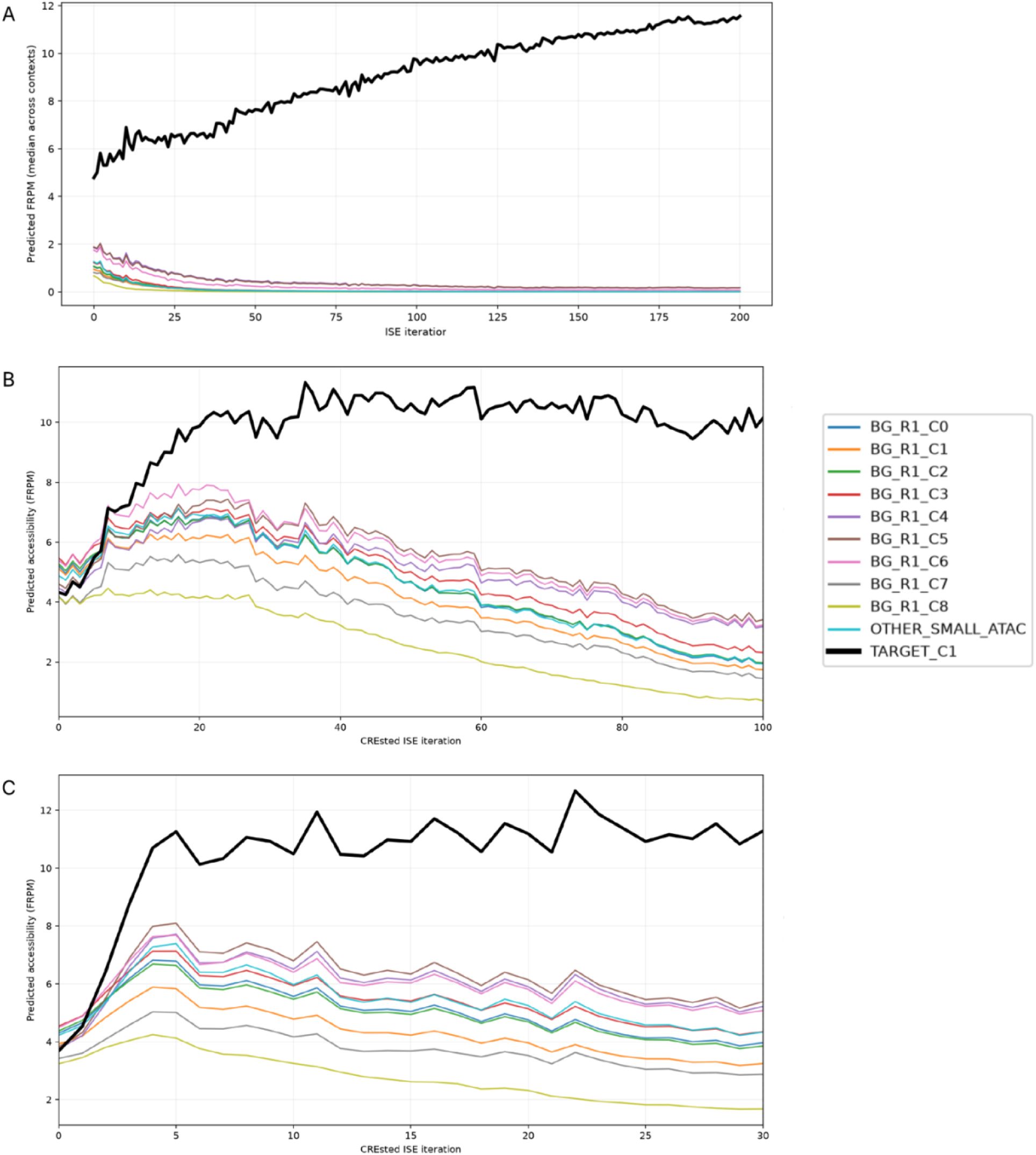
Sequence optimization increases predicted accessibility separation between C1 and comparator populations. Example trajectories of Borzoi-predicted accessibility, expressed as normalized ATAC fragment counts during in silico evolution (ISE). **A** 1,000-bp sequence optimized for 200 iterations. **B** A 500-bp sequence optimized for 100 iterations. **C** A 200-bp sequence optimized for 30 iterations.

The 200-bp library included 1,024 inserts, comprising 512 natural and 512 random starting sequences, optimized over 30 in silico evolution (ISE) iterations. The median C1-to-strongest-competitor ratio increased 1.74-fold for natural starts and 2.21-fold for random starts (Table 1). Zero optimized 200-bp sequences achieved a C1-to-competitor prediction ratio of at least 2 across all eight DNA backgrounds (Table 1).

**Table 1.**
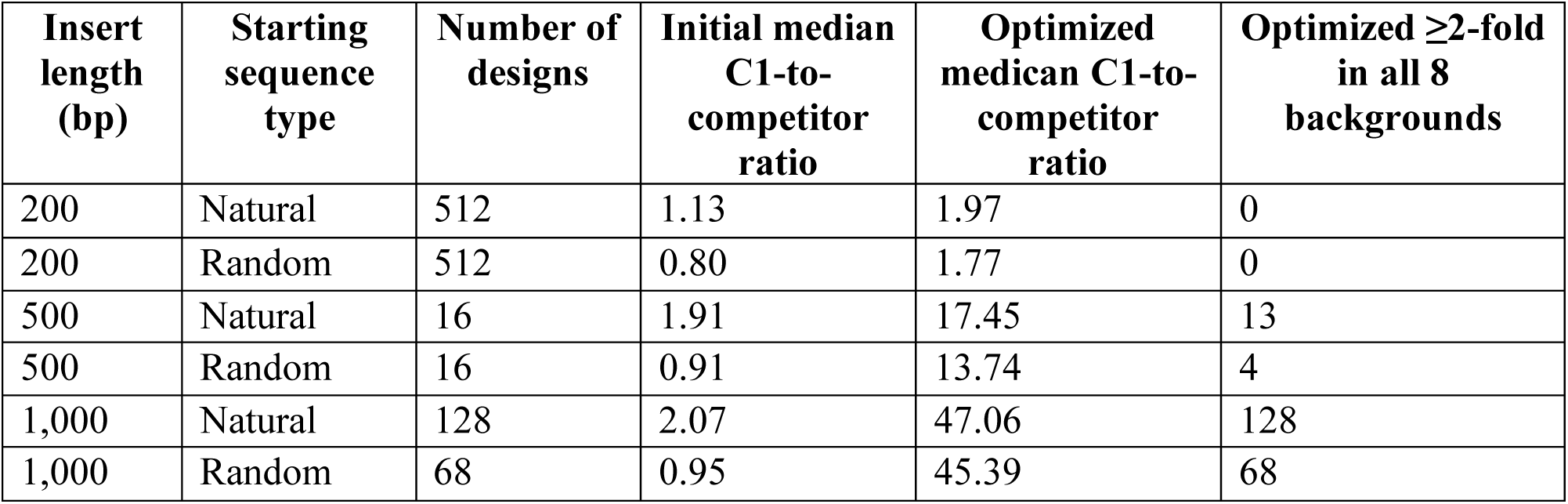
Predicted C1-selectivity prior to and after sequence optimization. Median C1-to-competitor ratio was calculated as the ratio of predicted normalized ATAC fragment counts for C1 divided by the highest predicted non-C1 population across all 8 DNA backgrounds. The optimized ≥2-fold in all 8 backgrounds column represents the number of optimized designs with predicted C1 counts ≥2x the highest non-C1 prediction in each of the 8 DNA backgrounds.

| **Insert length (bp)** | **Starting sequence type** | **Number of designs** | **Initial median C1-to-competitor ratio** | **Optimized medican C1-to-competitor ratio** | **Optimized ≥2-fold in all 8 backgrounds** |
|---|---|---|---|---|---|
| 200 | Natural | 512 | 1.13 | 1.97 | 0 |
| 200 | Random | 512 | 0.80 | 1.77 | 0 |
| 500 | Natural | 16 | 1.91 | 17.45 | 13 |
| 500 | Random | 16 | 0.91 | 13.74 | 4 |
| 1,000 | Natural | 128 | 2.07 | 47.06 | 128 |
| 1,000 | Random | 68 | 0.95 | 45.39 | 68 |

The 500-bp pilot included 32 inserts, including 16 natural and 16 random starts, optimized over 100 ISE iterations. The median C1-to-strongest-competitor ratio increased 9.14-fold for natural starts and 15.10-fold for random starts (Table 1). 13 natural-start and 4 random-start optimized sequences achieved at least a 2-fold predicted C1 preference in all 8 DNA backgrounds (Table 1).

Lastly, 128 natural-start and 68 random-start 1,000-bp inserts were optimized over 200 ISE iterations. Natural starts achieved a 22.73-fold improvement in the median C1-to-strongest-competitor ratio, and random starts achieved a 47.78-fold improvement (Table 1). All 196 retained 1,000-bp designs achieved at least a 2-fold predicted C1 preference in all 8 DNA backgrounds (Table 1). These runs differed in insert length, iteration count, starting sequences, and sample size, so they do not isolate the effect of insert length.

### Optimized sequences contain enriched sequence patterns

STREME identified five sequence patterns enriched in optimized sequences compared to their corresponding starting sequences and six sequence patterns enriched in optimized sequences compared to dinucleotide-matched controls. (E<0.05) (Figure 5). Eight motifs were able to be matched to JASPAR 2026 reference profiles with Tomtom (q≤0.05), while three had no significant matches (Table 2).

**Figure 5.**
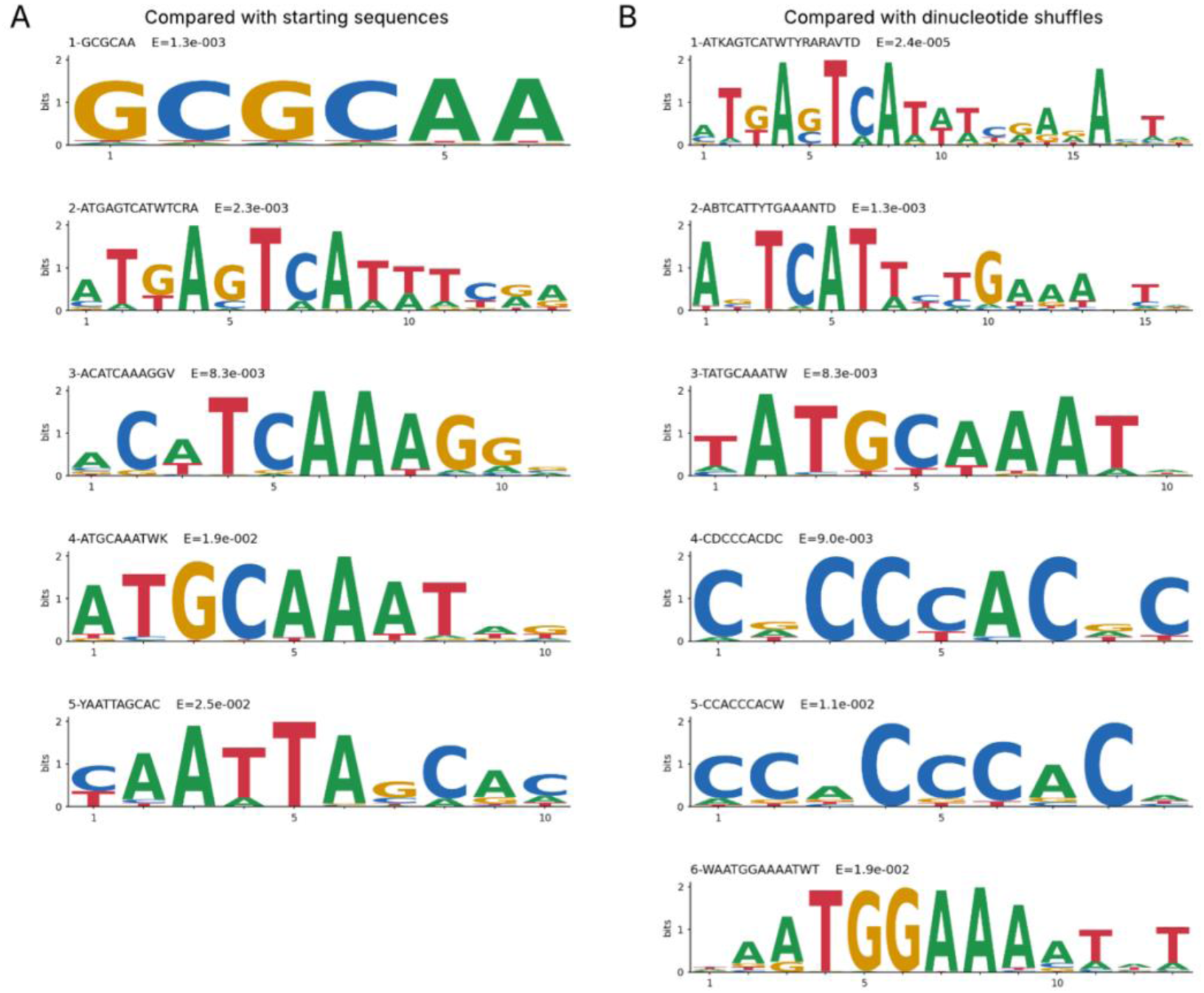
STREME identifies enriched sequence patterns in optimized 1,000-bp sequences. Patterns with E < 0.05 are shown, with E-values displayed above each logo. **A** Sequence logos of patterns enriched relative to starting sequences. **B** Sequence logos of patterns enriched relative to dinucleotide-shuffled controls.

**Table 2.**
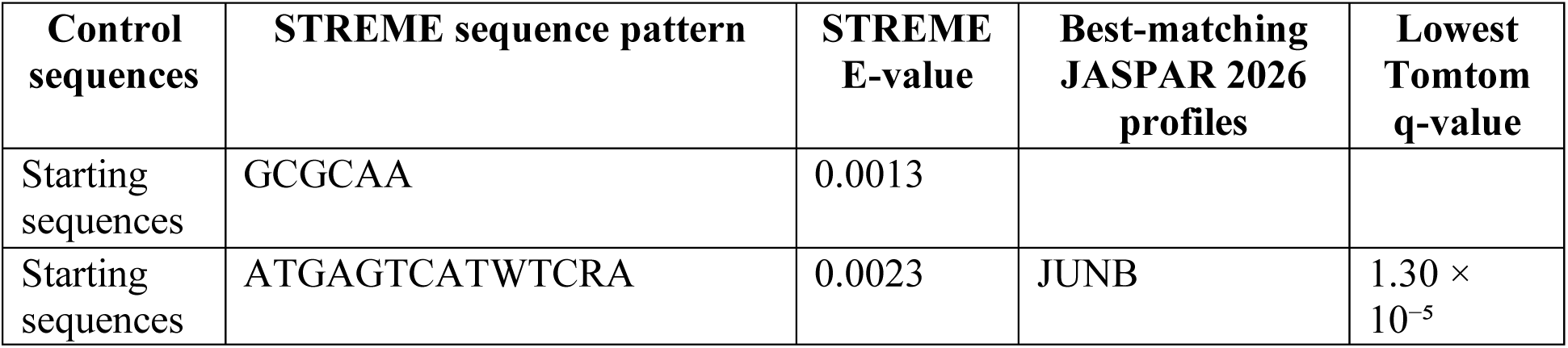
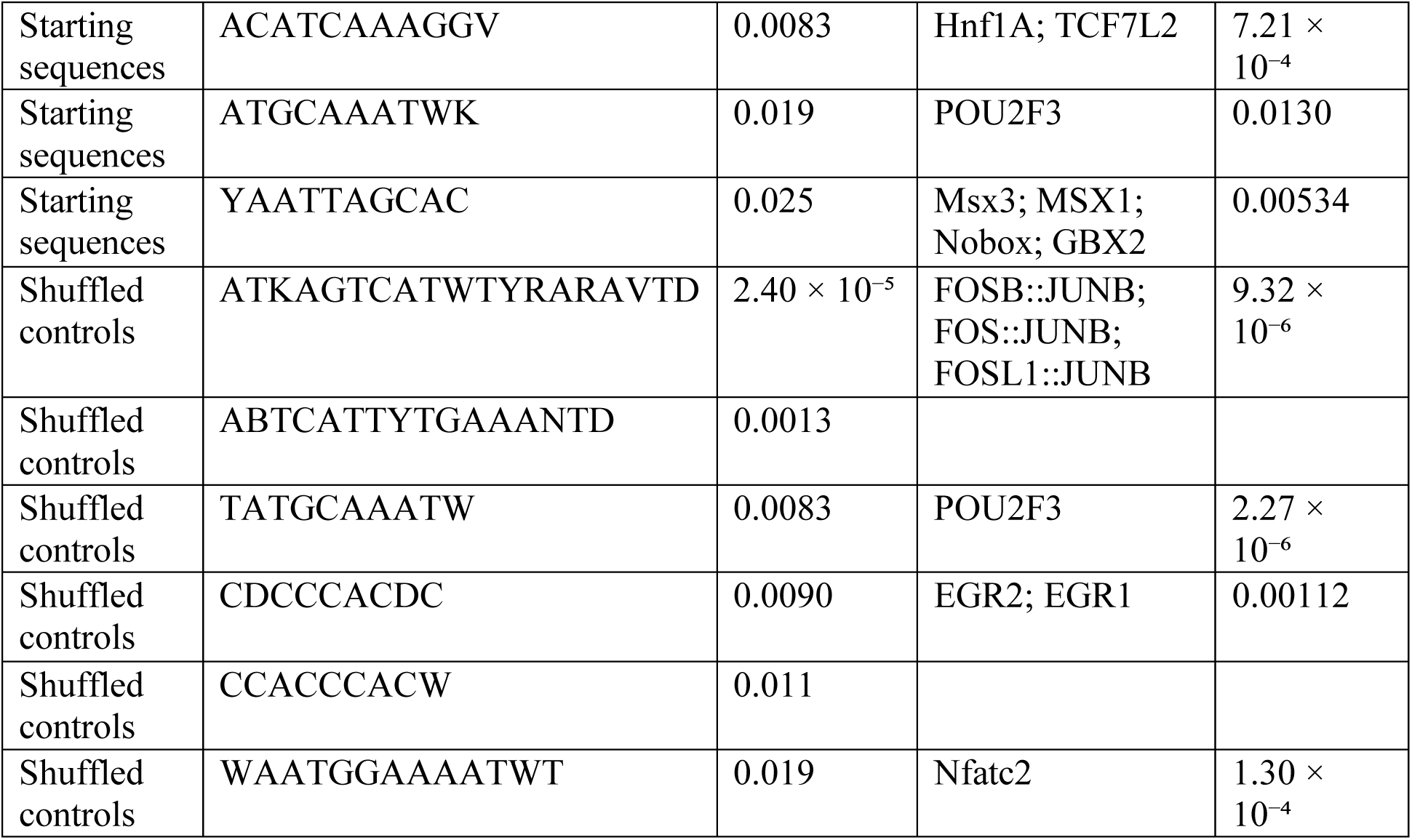
Enriched STREME sequence patterns and their JASPAR 2026 reference matches. E-values describe enrichment relative to the indicated controls. Tomtom q-values describe similarity to reference motifs. Multiple JASPAR profiles are listed when the best q-value is tied.

Shared JASPAR matches between optimized sequences compared with starting sequences and with dinucleotide shuffles include JUN/FOS-related and POU reference profiles (Table 2).

### Known motif scanning identifies increased POU-, NFAT-, and TCF/LEF-related sequence matches

The FIMO scan identified changes in motif occurrences in optimized sequences compared with initial sequences (Table 3). Matches to POU, NFAT, and TCF/LEF JASPAR profiles were more frequent in optimized sequences.

**Table 3.** FIMO JASPAR matches before and after sequence optimization. Values indicate sequences containing at least one FIMO match. The top 10 largest increases in sequence presence are shown.

| **JASPAR 2026 profile** | **Starting sequences with ≥1 match (n/196)** | **Optimized sequences with ≥1 match (n/196)** |
|---|---|---|
| POU2F3 | 60 | 165 |
| NFATC4 | 47 | 151 |
| Hnf1A | 60 | 165 |
| TCF7L2 | 64 | 167 |
| Pou5f1::Sox2 | 74 | 168 |
| Lef1 | 48 | 152 |
| Nfatc2 | 79 | 166 |
| POU2F1 | 59 | 157 |
| POU5F1 | 52 | 154 |
| POU5F1B | 63 | 159 |

### IG and EIG recover shared AP-1-associated sequence patterns in natural and optimized C1 regions

IG analysis of natural C1 sequences identified 15 positive and 6 negative sequence patterns, while EIG identified 13 positive and 7 negative patterns (Figures 6A and 6C). Both IG and EIG recovered dominant AP-1-associated patterns matching FOS::JUN (2,406 IG seqlets) and FOSL2::JUN (2,114 EIG seqlets), respectively. Both methods also recovered patterns matching IRF1 (266 IG and 202 EIG seqlets), ETV2::FOXI1 (130 IG and 98 EIG seqlets), and POU2F3 (61 IG and 55 EIG seqlets), along with additional patterns matching NFAT-, TCF7-, and ZNF-family motifs. IG additionally identified patterns matching RUNX1, NFKB2, CREB3L4, CREM, and JUND, while EIG identified patterns matching FOSL2::JUND and RELA.

**Figure 6.**
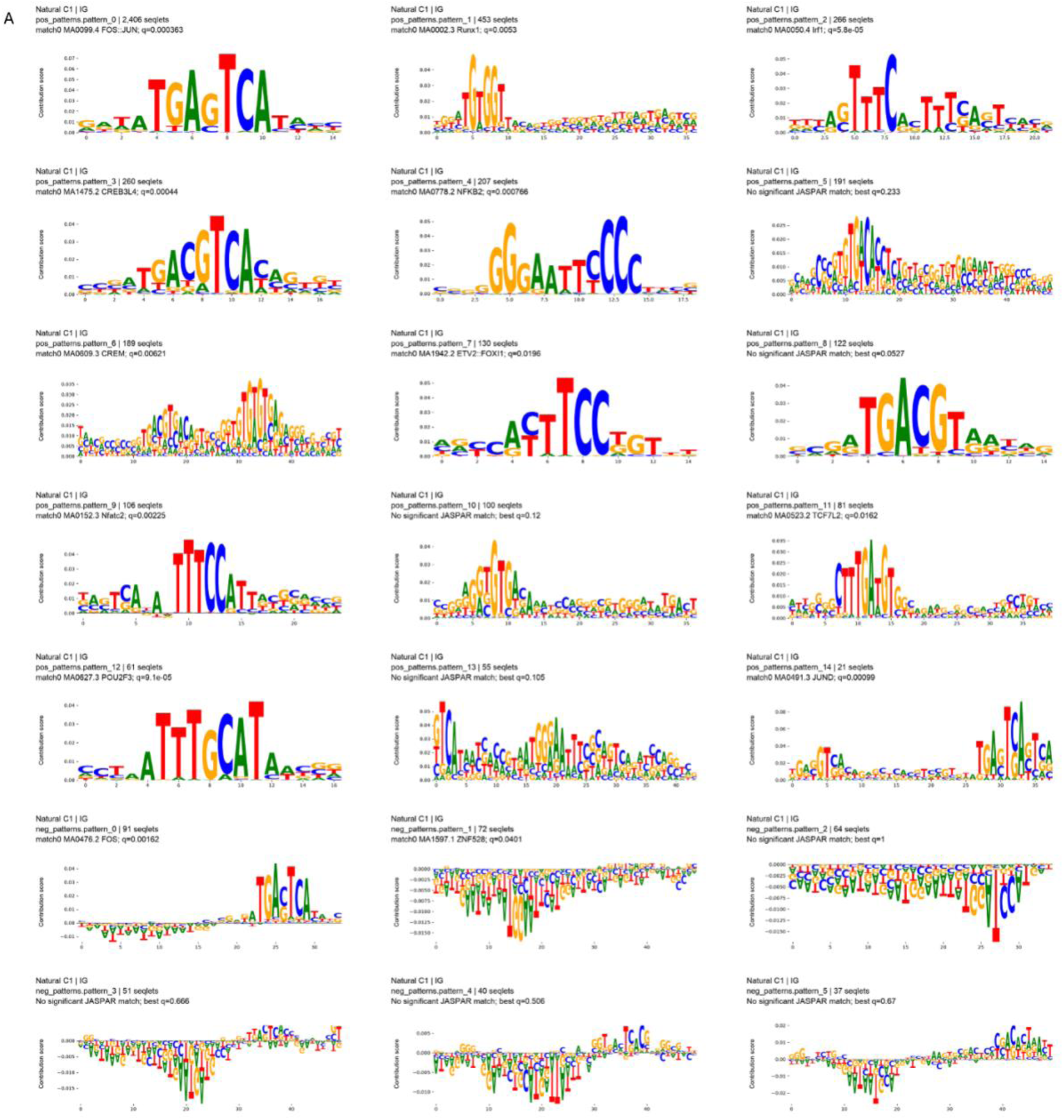
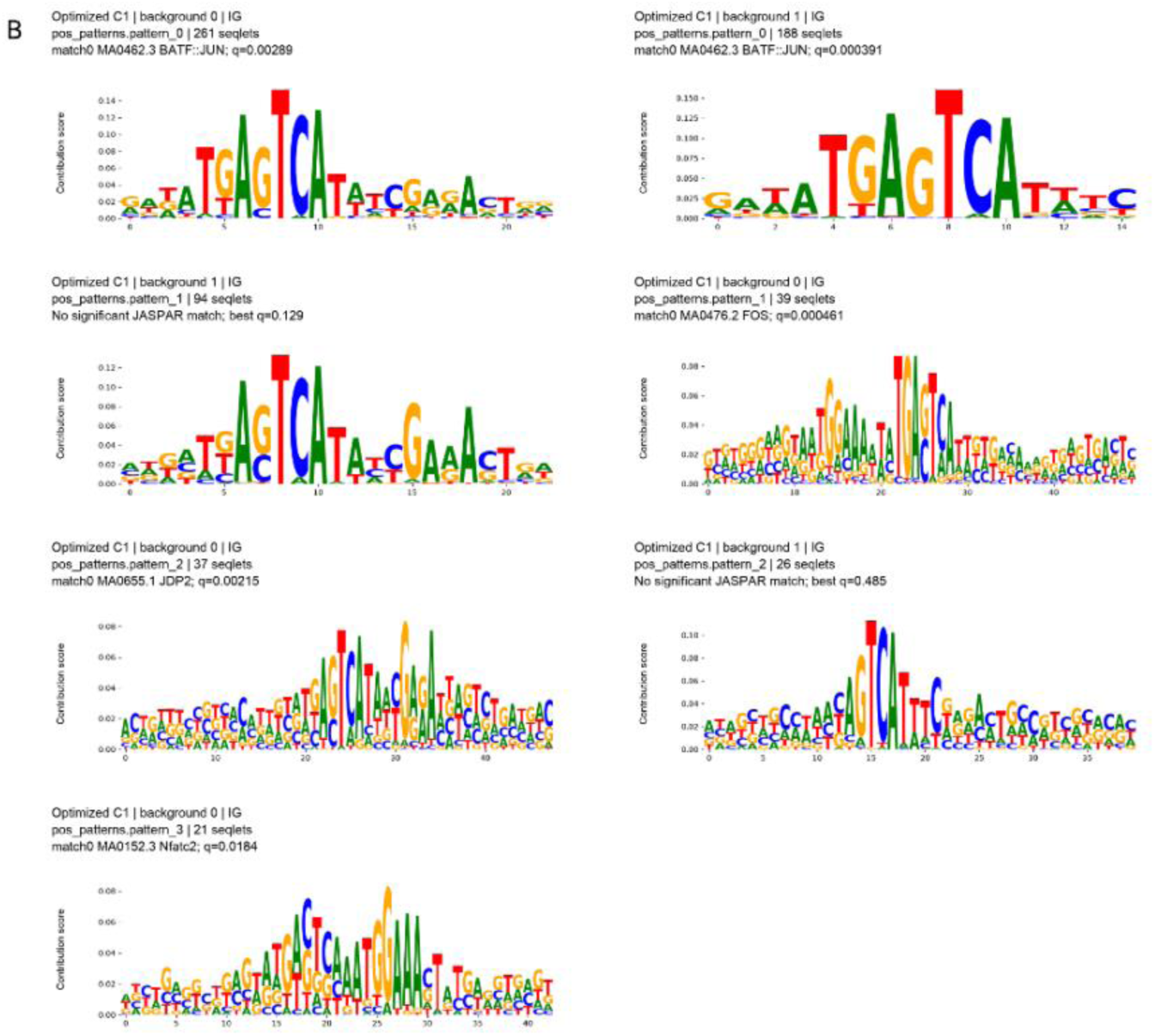
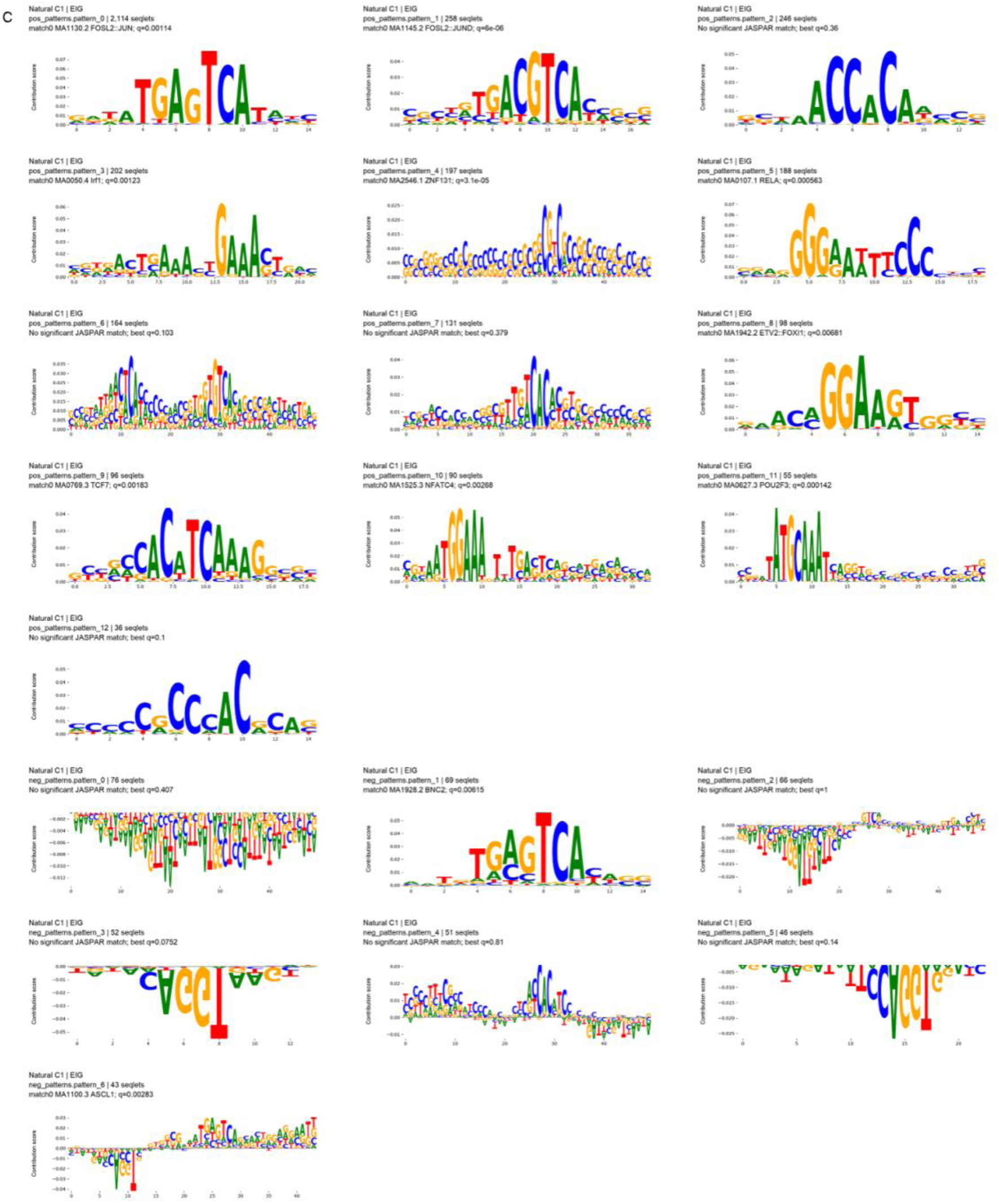
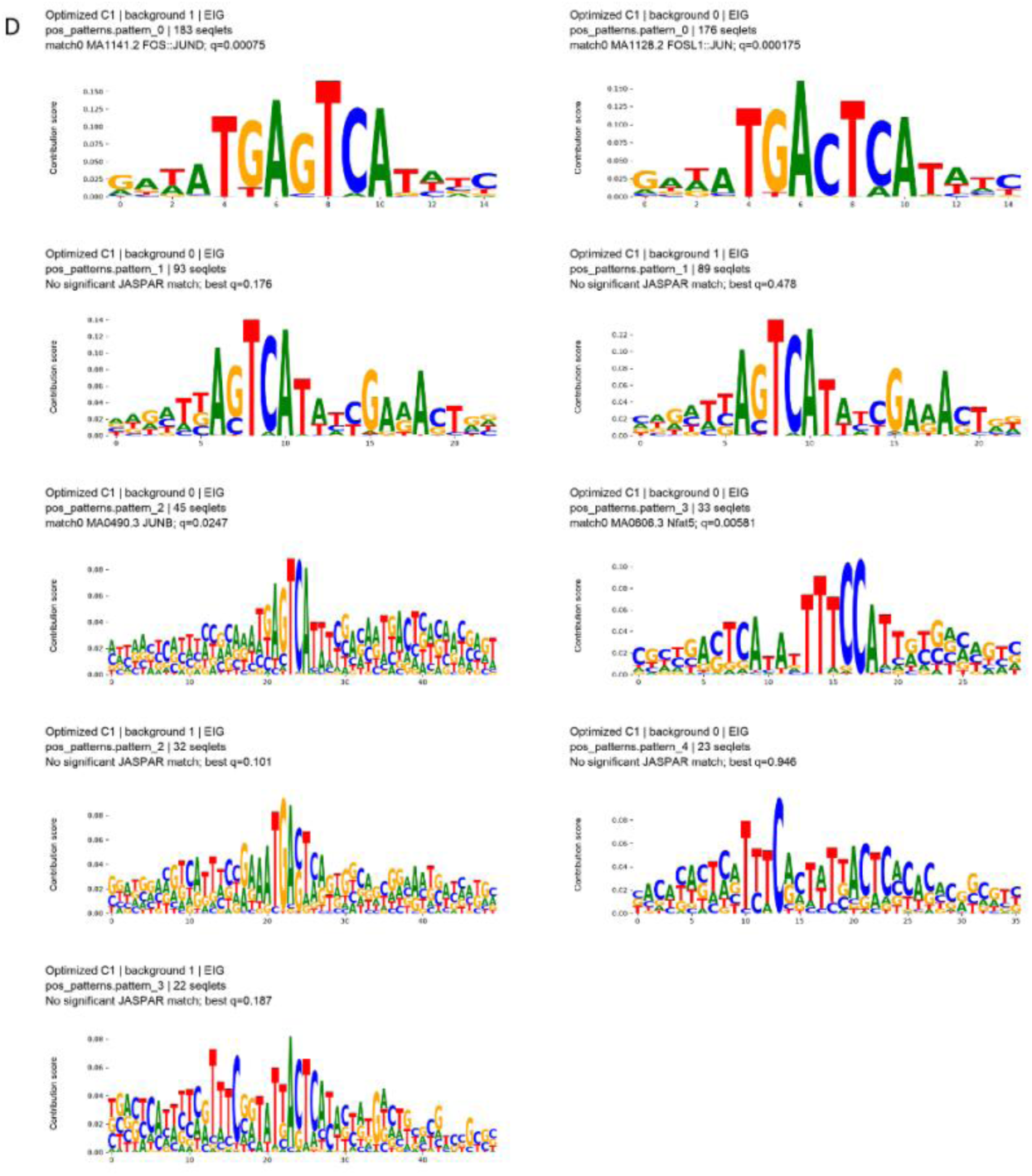
TF-MoDISco analysis of IG and EIG identifies AP-1-associated sequence patterns in natural and optimized C1 regions. Sequences represent positive and negative contribution patterns. Patterns are annotated with the number of TF-MoDISco-identified seqlets, the most significant JASPAR 2026 match, and the q-value of the JASPAR match. **A** IG patterns from natural sequences. **B** IG patterns from optimized sequences. **C**. EIG patterns from natural sequences. **D** EIG patterns from optimized sequences.

In optimized sequences, IG identified seven positive patterns and 0 negative patterns, while EIG identified nine positive patterns and 0 negative patterns (Figures 6B and 6D). IG recovered BATF::JUN-matching patterns in backgrounds 0 and 1, containing 261 and 188 seqlets, respectively. IG also identified patterns matching FOS, JDP2, and NFATC2, while EIG identified patterns matching FOS::JUND, FOSL1::JUN, JUNB, and NFAT5.

## Discussion

Here, we applied CREsted to design candidate synthetic CREs predicted to be preferentially accessible in an ATAC-defined Tex-prog-like CD8+ T-cell population, C1. Targeting Tex-prog-like cells could enable cell-state-specific transgene expression, but distinguishing these cells from other Tex cells is a challenge due to their many shared regulatory features. Using the fine-tuned Borzoi model, CREsted generated optimized sequences with substantially higher predicted accessibility in C1. These findings suggest that sequence optimization can identify candidate CREs that may distinguish Tex-prog-like cells from closely related CD8+ T-cell states.

Evaluation of recurring patterns in optimized sequences identified AP-1-, NFAT-, POU-, EGR-, and TCF/LEF-associated patterns. Furthermore, contribution scores analysis identified similar AP-1, BATF-, NFAT-, POU-, IRF-, and TCF/LEF-associated patterns. These recovered sequence patterns are broadly consistent with known regulatory programs associated with T-cell exhaustion. Walker et al. found that Tex-prog-like CD8+ T-cell states were enriched for transcription factor motifs associated with both follicular helper CD4+ T cells and exhausted CD8+ T cells, including AP-1, LEF1, TCF1, POU2F2, NFAT family, RUNX, and T-bet motifs.^16^ Because IG and EIG explain predicted C1 counts rather than the C1-to-competitor objective, these patterns do not establish which motifs confer C1 selectivity. An EGR-family sequence enriched relative to dinucleotide-shuffled controls matched both EGR2 and EGR1 reference profiles (Table 2). Wagle et al. showed that EGR2 is selectively expressed in progenitor-exhausted T cells and reduced in more differentiated states.^17^ The motif match alone does not establish EGR2 binding.

However, several limitations should be considered. C1 was identified through ATAC-based clustering and represents a Tex-prog-like population, not a purified Tex-prog population. Since the analysis combined multiple studies across varying disease contexts, differences between populations may reflect experimental context rather than exhaustion state alone. Furthermore, the same Borzoi model was used to optimize and evaluate the designed sequences and does not provide independent validation. The optimization objective could improve relative selectivity by reducing predictions of competitor populations, meaning that high C1-to-competitor ratios do not necessarily indicate strong C1 activity. Additionally, predicted chromatin accessibility does not establish transcription factor binding or enhancer-driven transcription.

These results support the identification of candidate synthetic CREs with predicted preferential accessibility in C1, but their functional specificity remains to be experimentally evaluated. Future studies will evaluate these optimized sequences alongside dinucleotide-matched controls in primary mouse CD8+ T cells following LCMV Clone 13 infection. These experiments will determine whether the predicted C1 preference translates into functional enhancer activity and cell-state specificity.

## Methods

### Data sources

Publicly available paired single-cell RNA-seq (scRNA-seq) and single-cell ATAC-seq (scATAC-seq) data from mouse infection and tumor models were obtained from four studies deposited in the Gene Expression Omnibus (GEO): GSE230363, GSE267584, GSE255042, GSE290618.^15,18,19,20^ From GSE230363, we included naïve P14 CD8+ T cells and cells on days 8, 15, and 30 after infection with either LCMV Armstrong or LCMV Clone 13. From GSE267584, we included naïve CD8+ T cells, splenic and small intestinal intraepithelial lymphocyte (IEL) cells collected at day 6 and 42 post-infection with LCMV Armstrong. Splenic cells at day 6 and 42 following infection with LCMV Clone 13, and cells from tumor models mWnt-GP, KPC-GP, B16-GP, and splenic cells from B-ALL-GP 7 days post infection were included. From GSE255042, we included naïve splenic P14 CD8+ T cells, and from GSE290618, we included T cells from tumor and adjacent tumor-free cecal tissues collected 4 weeks after tumor transplantation.

### RNA count extraction and quality control

scRNA-seq count matrices were downloaded from GEO and processed using Python and Scanpy (1.10.3).^21^ Gene expression features were retained. Cells were restricted to CD8 populations. Gene symbols were standardized and counts from features sharing a canonical symbol were summed. Cells with ≤ 500 RNA counts, >20% mitochondrial RNA, and ≤300 detected genes were removed from analysis. Doublets were assessed for each dataset using Scrublet (30 pc, doublet rate .05) and DoubletDetection(25 iterations, up to 10000 variable genes, 30 pc, boost rate 0.25, Louvain resolution 4, standard scaling, pseudocount 0.1).^22,23^ Suspected doublets were excluded using a DoubletDetection threshold of p = 10^-7 and a voting fraction of ≥0.9. For GSE290618, doublet modeling used the entire cell population. In reviewing doublet populations, we retained GSE230363 GSM7220515 cells in a cycling cluster with Scrublet values ≤0.30.

### Reference annotations

Quality-controlled, unnormalized RNA counts were submitted to the online ImmGenT T-RBI CD8 portal for annotation. ImmGenT T-RBI returned fine annotations for 160,955 cells. 818 cells were filtered by the T-RBI.

Separately cells from GSE267584 were processed and labeled according to Green et al. labeling 26 paper-associated states. All cells were mapped to a GSE267584 reference containing 26 cell states using Seurat RNA anchor-based label transfer (LogNormalize, scale factor = 10,000, up to 2,000 shared variable genes, 30 PCA dimensions, seed = 1729, reduction = pcproject, k.anchor = 5, k.weight ≤ 50).^24^

### RNA-ATAC linkage, peak calling and ATAC QC

scRNA-seq and scATAC-seq data were matched using source-aware cell identifiers and multiome barcodes. scATAC-seq fragments were restricted to primary nuclear mm10 chromosomes. ATAC peaks were called using MACS2 (BEDPE mode, mouse genome size, q-value threshold of 0.05) on pooled fragments from all candidate CD8 cells within each experimental capture and, separately, from ImmGenT-annotated groups containing at least 50 cells within each capture.^24^ Peaks from 199 pseudobulks were combined into a consensus set of 500-bp regions, excluding mm10 v2 blacklist regions, with pycisTopic.^25,26^ This resulted in 439,167 consensus peak regions.

TSS enrichment was calculated with GENCODE M25 protein-coding transcription start sites.^27^ Cells with <1,000 unique primary nuclear fragments, TSS enrichment <5, and a fraction of fragments in consensus peaks <0.20 were excluded. 117,653 paired cells were retained. Regions detected in <10 cells were removed, retaining 439,167 regions.

### scRNA-seq and scATAC-seq integration

RNA counts were normalized to 10,000 counts per cell and log transformed. RNA dimensionality reduction was performed (n_top_genes=3000, flavor=Seurat, batch_key= source study). Mitochondrial, ribosomal, and TCR variable and joining genes were excluded.

Expression was scaled (max_value=10) and 50 principal components were calculated. Batch correction was performed on the first 30 PCs using Harmony (source study as batch, maximum 20 iterations), followed by UMAP (n_neighbors=30, minimum distance = 0.3).^28,29^

ATAC dimensionality reduction was performed using latent semantic indexing (LSI) on binarized accessibility counts for peaks detected in ≥30 cells. TF-IDF transformation (L2 normalization, smooth IDF, IDF weighting, no sublinear term-frequency transformation) was followed by truncated singular value decomposition (51 components, 5 iterations). The first component was excluded because of its association with sequencing depth. The retained 50 components were standardized and batch-corrected with Harmony (maximum iterations = 20). Cells expressing Cd8a or Cd8b were retained, yielding a final dataset of 111,347 cells.

### ATAC cluster definitions

A 30-nearest-neighbor graph was constructed using Harmony-corrected LSI components. Leiden clustering was performed (resolution=0.3, 0.5, 1.0, iterations=2).^30^ Resolution 0.3 identified 4 clusters, including C1, which contained 4,440 cells, and defined our primary target. Resolution 1.0 clusters provided finer comparator populations for the Borzoi model. Most cells in resolution 0.3 cluster C1 (4,164 of 4,440; 93.8%) were assigned to cluster C5 at resolution 1.0. C5 also contained cells outside C1, which were used as a closely related comparator for RNA expression analyses.

### C1 RNA characterization

C1 was characterized by the proportion of cells assigned to each ImmGenT and Green et al. reference annotation. Marker-gene expression was examined for C1 compared with non-C1 cells using normalized RNA expression.

### Accessibility targets

ATAC fragments overlapping each 500-bp peak were summed for each population. Counts were normalized per million unique fragments within each population. CREsted’s normalize_peaks function (1.10.0) (top_k_percent=0.03) was used to estimate scaling factors from training peaks, which were applied to all peaks.

### Borzoi fine-tuning

A Borzoi model was fine-tuned to predict normalized ATAC fragment counts for C1 and 10 comparator populations.^31^ Model inputs were 2,048-bp mm10 DNA sequences centered on the 500-bp peaks. The chromosome split contained 356,938 training peaks, 45,032 validation peaks from chromosomes 8 and 10, and 37,197 test peaks from chromosomes 9 and 18. The chromosome partition appled during fine-tuning was not matched to the original Borzoi pretraining folds. The exclusion of the validation and test regions from pretraining was not established. The model had 11 outputs: C1, nine resolution-1.0 ATAC groups containing at least 200 non-C1 cells, and a pooled output for smaller groups. C1 cells were excluded from comparator populations.

Borzoi fine-tuning was initialized from the pretrained mouse checkpoint borzoi_mouse_rep0. Following the CREsted tutorial, the original prediction head was dropped and replaced by a Flatten layer followed by a new dense layer (activation=softplus) after final_conv_activation. All layers were fine-tuned. Training used a batch size of 32, CosineMSELogLoss(max_weight=100, multiplier=1), Adam with an initial learning rate (LR) of 10⁻⁵, epsilon of 10⁻⁷, gradient-norm clipping at 5, FP32 precision, and reverse-complement (RC) augmentation.^32^

The LR was halved after three epochs without improvement in the validation macro-averaged average precision for population-selective peaks, with a minimum learning rate of 10^-6. The training allowed 20 epochs with early stopping after 7 epochs without improvement; early stopping was triggered after epoch 15.

Validation macro average precision (AP) was calculated across populations with ≥200 cells, ≥50 selective training peaks and ≥10 selective validation peaks. Resolution-1.0 ATAC clusters C4, C6, C7, C8 and C1 met these criteria. Macro-AP was calculated as the unweighted mean of each AP value. All 11 population outputs were used for model training.

Selective peaks required ≥10 raw ATAC fragments aggregated across cells in the target population and measured normalized fragment counts ≥2 times those of the strongest comparator population. The pooled small population was excluded from the comparator set for both validation and primary test evaluation. For C1, positive peaks were defined relative to the nine non-C1 resolution-1.0 ATAC populations. For each other evaluated population, comparators comprised the remaining individually modeled populations, including C1, with the pooled small population excluded.

Forward and RC predictions were averaged, and population-specific scaling factors were removed prior to evaluation. Validation and test regions were ranked with the same scoring rule: the difference between log1p-transformed predicted normalized fragment counts for the target population and its strongest comparator population. The pooled small population was excluded from this ranking score in both validation and primary test evaluation.

Training completed after 15 epochs, with epoch 8 achieving the highest validation macro-averaged AP for selective peaks. Model performance was evaluated on chromosomes held out from fine-tuning using AP and precision at the top 20, 50, 100, and 500 predictions for C1-selective peaks. A separate test sensitivity analysis included the pooled small population in both the positive-peak definition and the ranking score.

### Sequence optimization

Synthetic sequence optimization used CREsted’s in silico evolution (ISE) function (in_silico_evolution). Sequences were optimized using the frozen epoch-8 Borzoi checkpoint. Optimization used 30 iterations for 200-bp sequences, 100 iterations for 500-bp sequences, and 200 iterations for 1,000-bp sequences. No early stopping or known motifs were applied.

The 200-bp run included 512 natural starting sequences from 50 training peaks and 512 composition-matched random starts. The 500-bp optimization was conducted as a pilot with 16 natural and 16 random starting sequences. The 1,000-bp ISE was planned to include 128 natural and 128 random starts. Sequence generation was stopped early at 128 natural and 68 random starts.

Within each planned natural-start and random-start group, an equal number of designs were randomly assigned a C1-predicted retention requirement of either 25% or 50% of their starting C1 prediction. The minimum C1 value was defined as the greater value of either the C1-prediction retention requirement or 25% of the median natural-start C1 prediction. ISE predictions that fell below the minimum C1 requirement incurred a penalty (10 * the difference between the log1p-transformed minimum requirement and the C1 prediction averaged across the two optimization backgrounds).

Candidates were inserted at the center of two fixed 2,048-bp DNA backgrounds from training chromosomes. For each background, the selectivity score was the difference between log1p-transformed predicted fragment counts for C1 and the strongest competing population among all 10 non-C1 outputs, including the pooled small population. The optimization objective was defined by the average of the average and minimum of the two backgrounds minus the minimum C1 requirement penalty if applicable. This objective was chosen to favor high performance in both backgrounds while allowing the predicted C1 activity to decrease as long as specificity increased.

After optimization, the highest-scoring sequences meeting the C1 minimum in both backgrounds were retained. All designs were evaluated in an additional 8 audit backgrounds from training chromosomes. Forward and RC predictions were averaged and population scaling factors were removed. The strongest competitor was the maximum prediction among all 10 non-C1 populations. For each design, the C1-to-competitor ratio was calculated across all 8 audit backgrounds.

### 1,000-bp Motif discovery with STREME and FIMO

STREME (MEME Suite 5.5.9) was used to identify enriched 6-20-bp sequence patterns in the 196 optimized 1,000-bp sequences.^33^ Optimized sequences were compared with their corresponding starting sequences and, separately, with shuffled, dinucleotide frequency-matched copies of optimized sequences. STREME used 20 output motifs per comparison, a first-order background model, and a holdout fraction of 0.1. 177 sequences were used for discovery and 19 were used for internal significance estimation. Sequence patterns with reported E-values < 0.05 were retained.

STREME motif probability matrices were compared with the 1,019 JASPAR 2026 CORE vertebrate nonredundant profiles using Tomtom (MEME Suite 5.5.9).^34^ Matches with q≤0.05 were retained.

FIMO (MEME Suite 5.5.9) was used to scan starting and optimized 1,000-bp sequences for matches to JASPAR 2026 CORE vertebrate nonredundant profiles (NRDB nucleotide background, p-val <.0001).^35,36^ Motif presence was defined as at least one passing match per sequence. Changes in motif presence were summarized descriptively.

### Natural region selection

For each of the 11 Borzoi model populations, we selected the top 2,000 population-specific natural regions for comparison. For each region and population, normalized ATAC fragment counts and model predictions were placed on the same CREsted-scaled units and averaged. CREsted’s sort_and_filter_regions_on_specificity (method= ‘gini’, top_k=2,000) was used to select the natural regions from all chromosomes. No minimum Gini-score was applied. The C0 population contained 480 positive-score entries and 1,520 zero-score entries.

### Contribution score calculation

Contribution scores were calculated using the frozen epoch-8 Borzoi checkpoint. Forward and RC predictions were averaged and population scaling factors were removed. CREsted’s contribution_scores function was used to calculate integrated gradients (IG) using a zero baseline and 26 integration points.^37^ Expected integrated gradients (EIG) were calculated with 25 randomly permuted versions of the input as baselines and 26 integration points per baseline.^38^ The integration batch size was 4 and simplex correction was not used. Attribution was performed for population-specific predicted counts. Contribution scores were calculated for both the 22,000 natural regions (2,000 regions per 11 output groups) and 784 optimized stages (196 starting and optimized sequences in both backgrounds).

### TF-MoDISco pattern discovery and pattern grouping

TF-MoDISco (tfmodisco-lite 3.0.0) pattern discovery was performed separately for IG and EIG using CREsted.^39^ Discovery was performed across natural and synthetic sequences. TF-MoDISco was run on the central 1,000-bp sequences, with a 20-bp sliding window, 5-bp flanks, a maximum of 20,000 seqlets, a minimum metacluster size of 100, a minimum final cluster size of 20, two Leiden runs, and a target seqlet FDR of 0.05.

IG and EIG patterns were grouped separately for natural and synthetic sequences using CREsted’s process_patterns function (method=’greedy’, similarity_threshold=6, trim_threshold=0.025, discard_threshold=0.1). Tomtom (q≤0.05) was used to compare sequence patterns to the JASPAR 2026 CORE vertebrate profiles.

## Data and code availability

Publicly available sequencing scRNA-seq and scATAC-seq data analyzed in this study are deposited in GEO under accessions GSE230363, GSE267584, GSE255042 and GSE290618. Starting and candidate sequences for the 200-bp, 500-bp and 1,000-bp libraries are provided in Supplementary File 1, and saved dinucleotide-shuffled audit controls are provided in Supplementary File 2. DNA backgrounds, per-design predictions, population labels, chromosome-split assignments, random seeds, supporting analysis outputs and the trained C1 Borzoi checkpoint are available in Zenodo at 10.5281/zenodo.24480177. Custom code and configuration files for data preprocessing, RNA/ATAC integration, Borzoi model fine-tuning, sequence optimization and motif analysis are available at https://github.com/nazbuhn/Tex-prog-like_synthetic_cre_design. A snapshot of the code is archived in the same Zenodo record.

## References

1. Pauken, K. E. et al. Epigenetic stability of exhausted T cells limits durability of reinvigoration by PD-1 blockade. Science 354, 1160–1165 (2016). 10.1126/science.aaf2807

2. Masopust, D. et al. Guidelines for T cell nomenclature. Nat. Rev. Immunol. 26, 298–313 (2026). 10.1038/s41577-025-01238-2

3. Beltra, J.-C. et al. Developmental relationships of four exhausted CD8+ T cell subsets reveals underlying transcriptional and epigenetic landscape control mechanisms. Immunity 52, 825–841.e8 (2020). 10.1016/j.immuni.2020.04.014

4. Miller, B. C. et al. Subsets of exhausted CD8+ T cells differentially mediate tumor control and respond to checkpoint blockade. Nat. Immunol. 20, 326–336 (2019). 10.1038/s41590-019-0312-6

5. Siddiqui, I. et al. Intratumoral Tcf1+PD-1+CD8+ T cells with stem-like properties promote tumor control in response to vaccination and checkpoint blockade immunotherapy. Immunity 50, 195–211.e10 (2019). 10.1016/j.immuni.2018.12.021

6. Im, S. et al. Defining CD8+ T cells that provide the proliferative burst after PD-1 therapy. Nature 537, 417–421 (2016). 10.1038/nature19330

7. Lan, X. et al. Antitumor progenitor exhausted CD8+ T cells are sustained by TCR engagement. Nat. Immunol. 25, 1046–1058 (2024). 10.1038/s41590-024-01843-8

8. Gosai, S. J. et al. Machine-guided design of cell-type-targeting cis-regulatory elements. Nature 634, 1211–1220 (2024). 10.1038/s41586-024-08070-z

9. Zhang, Z. et al. SPECIFIC: A systematic framework for engineering cell state-responsive synthetic promoters reveals key regulators of T cell exhaustion. Quant. Biol. e97 (2025). 10.1002/qub2.97

10. Yuan, H. & Kelley, D. R. scBasset: sequence-based modeling of single-cell ATAC-seq using convolutional neural networks. Nat. Methods 19, 1088–1096 (2022). 10.1038/s41592-022-01562-8

11. Yin, C. et al. Iterative deep learning design of human enhancers exploits condensed sequence grammar to achieve cell-type specificity. Cell Syst. 16, 101302 (2025). 10.1016/j.cels.2025.101302

12. Kempynck, N. et al. CREsted: modeling genomic and synthetic cell-type-specific enhancers across tissues and species. Nat. Methods 23, 946–959 (2026). 10.1038/s41592-026-03057-2

13. Green, W. D. et al. Enhancer-driven gene regulatory networks reveal transcription factors governing T cell adaptation and differentiation in the tumor microenvironment. Immunity 58, 1725–1741.e9 (2025). 10.1016/j.immuni.2025.04.030

14. Immgen Magill, I., et al. immgenT: a comprehensive reference of convergent T-cell states in the mouse. bioRxiv 2026**.01.30.702892** (2026). 10.64898/2026.01.30.702892

15. Giles, J.R., Globig, A.M., Kaech, S.M. & Wherry, E.J. CD8+ T cells in the cancer-immunity cycle. Immunity 56, 2231–2253 (2023). 10.1016/j.immuni.2023.09.005

16. Walker, S.K., van der Veeken, J., Rudensky, A.Y. & Pritykin, Y. Single-cell multi-omics reveals archetypal regulatory programs shared across T cell subsets. Cell Rep. 45, 118156 (2026). 10.1016/j.celrep.2026.118156

17. Wagle, M.V., Vervoort, S.J., Kelly, M.J. et al. Antigen-driven EGR2 expression is required for exhausted CD8+ T cell stability and maintenance. Nat. Commun. 12, 2782 (2021). 10.1038/s41467-021-23044-9

18. Baxter, A. E. et al. The SWI/SNF chromatin remodeling complexes BAF and PBAF differentially regulate epigenetic transitions in exhausted CD8+ T cells. Immunity 56, 1320–1340.e10 (2023). 10.1016/j.immuni.2023.05.008

19. Huang, Y. J. et al. Continuous expression of TOX safeguards exhausted CD8 T cell epigenetic fate. Sci. Immunol. 10, eado3032 (2025). 10.1126/sciimmunol.ado3032

20. Huang, X. et al. Opposing functions of distinct regulatory T cell subsets in colorectal cancer. Immunity 59, 145–160.e9 (2026). 10.1016/j.immuni.2025.11.014

21. Wolf, F. A., Angerer, P. & Theis, F. J. SCANPY: large-scale single-cell gene expression data analysis. Genome Biol. 19, 15 (2018). 10.1186/s13059-017-1382-0.

22. Wolock, S. L., Lopez, R. & Klein, A. M. Scrublet: computational identification of cell doublets in single-cell transcriptomic data. Cell Syst. 8, 281–291.e9 (2019). 10.1016/j.cels.2018.11.005

23. Gayoso, A., Shor, J., Carr, A. J., Sharma, R. & Pe’er, D. DoubletDetection (Version v3.0). Zenodo (2020). 10.5281/zenodo.2678041

24. Zhang, Y. et al. Model-based analysis of ChIP-Seq (MACS). Genome Biol. 9, R137 (2008). 10.1186/gb-2008-9-9-r137

25. Bravo González-Blas, C., et al. SCENIC+: single-cell multiomic inference of enhancers and gene regulatory networks. Nat. Methods 20, 1355–1367 (2023). 10.1038/s41592-023-01938-4

26. Amemiya, H. M., Kundaje, A. & Boyle, A. P. The ENCODE blacklist: identification of problematic regions of the genome. Sci. Rep. 9, 9354 (2019). 10.1038/s41598-019-45839-z

27. Frankish, A., et al. GENCODE 2021. Nucleic Acids Res. 49, D916–D923 (2021). 10.1093/nar/gkaa1087

28. Korsunsky, I. et al. Fast, sensitive and accurate integration of single-cell data with Harmony. Nat. Methods 16, 1289–1296 (2019). 10.1038/s41592-019-0619-0

29. Becht, E. et al. Dimensionality reduction for visualizing single-cell data using UMAP. Nat. Biotechnol. 37, 38–44 (2019). 10.1038/nbt.4314

30. Traag, V. A., Waltman, L. & van Eck, N. J. From Louvain to Leiden: guaranteeing well-connected communities. Sci. Rep. 9, 5233 (2019). 10.1038/s41598-019-41695-z

31. Linder, J. et al. Predicting RNA-seq coverage from DNA sequence as a unifying model of gene regulation. Nat. Genet. 57, 949–961 (2025). 10.1038/s41588-024-02053-6

32. Kingma, D. P. & Ba, J. Adam: a method for stochastic optimization. In *International Conference on Learning Representations* (ICLR, 2015). https://arxiv.org/abs/1412.6980

33. Bailey, T. L. STREME: accurate and versatile sequence motif discovery. Bioinformatics 37, 2834–2840 (2021). 10.1093/bioinformatics/btab203

34. Gupta, S., Stamatoyannopoulos, J. A., Bailey, T. L. & Noble, W. S. Quantifying similarity between motifs. Genome Biol. 8, R24 (2007). 10.1186/gb-2007-8-2-r24

35. Grant, C. E., Bailey, T. L. & Noble, W. S. FIMO: scanning for occurrences of a given motif. Bioinformatics 27, 1017–1018 (2011). 10.1093/bioinformatics/btr064

36. Baydar Ovek, D., et al. JASPAR 2026: expansion of transcription factor binding profiles and integration of deep learning models. Nucleic Acids Res. 54, D184–D193 (2026). 10.1093/nar/gkaf1209

37. Sundararajan, M., Taly, A. & Yan, Q. Axiomatic attribution for deep networks. In Proceedings of the 34th International Conference on Machine Learning 70, 3319–3328 (PMLR, 2017).

38. Erion, G. et al. Improving performance of deep learning models with axiomatic attribution priors and expected gradients. *Nat*. Mach. Intell. 3, 620–631 (2021).

39. Shrikumar, A. et al. Technical note on transcription factor motif discovery from importance scores (TF-MoDISco) version 0.5.6.5. *arXiv* (2018). 10.48550/arXiv.1811.00416

